## Supplementary Figures for "Feed-forward stimulation of CAMK2 by the oncogenic pseudokinase PEAK1 generates a therapeutically ‘actionable’ signalling axis in triple negative breast cancer"

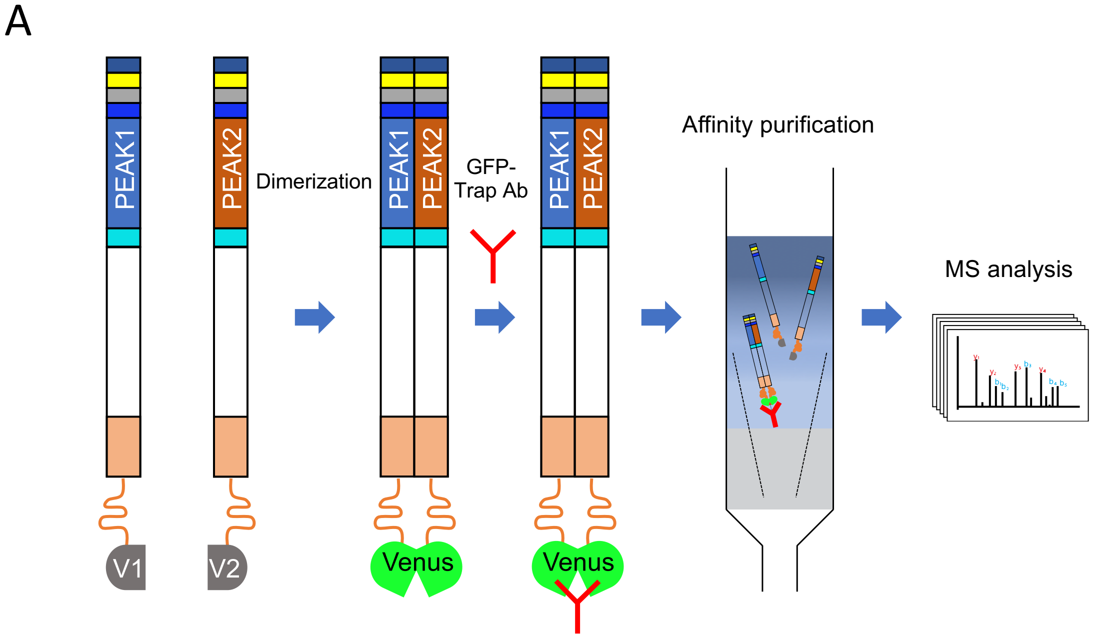


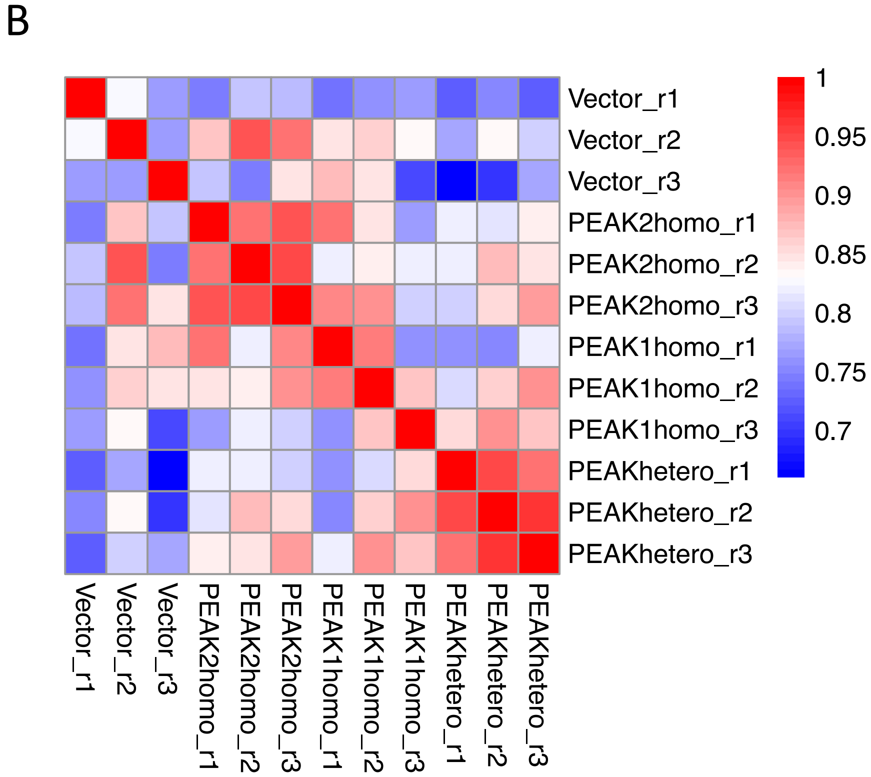


**Supp Figure 1. Characterization of PEAK complex interactomes via BiCAP-MS/MS. A. Schematic strategy of screen.** This is indicated for the PEAK1/PEAK2 heterodimer. PEAK1 and PEAK2 were fused to N-terminal (V1) and C-terminal (V2) fragments of the Venus protein, respectively. When PEAK1 and PEAK2 associate, V1 and V2 bind to each other and reconstitute Venus that can be recognized by the GFP-targeting nanobody (GFP-Trap). The heterodimer and its interactome can then be immunoaffinity purified for MS analysis. In the screen, this workflow was applied to the PEAK1 and PEAK2 homodimers as well as the heterodimer. **B. Sample correlation heatmap for the DIA MS analyses.** Data shown are for 3 replicates, r1-r3.

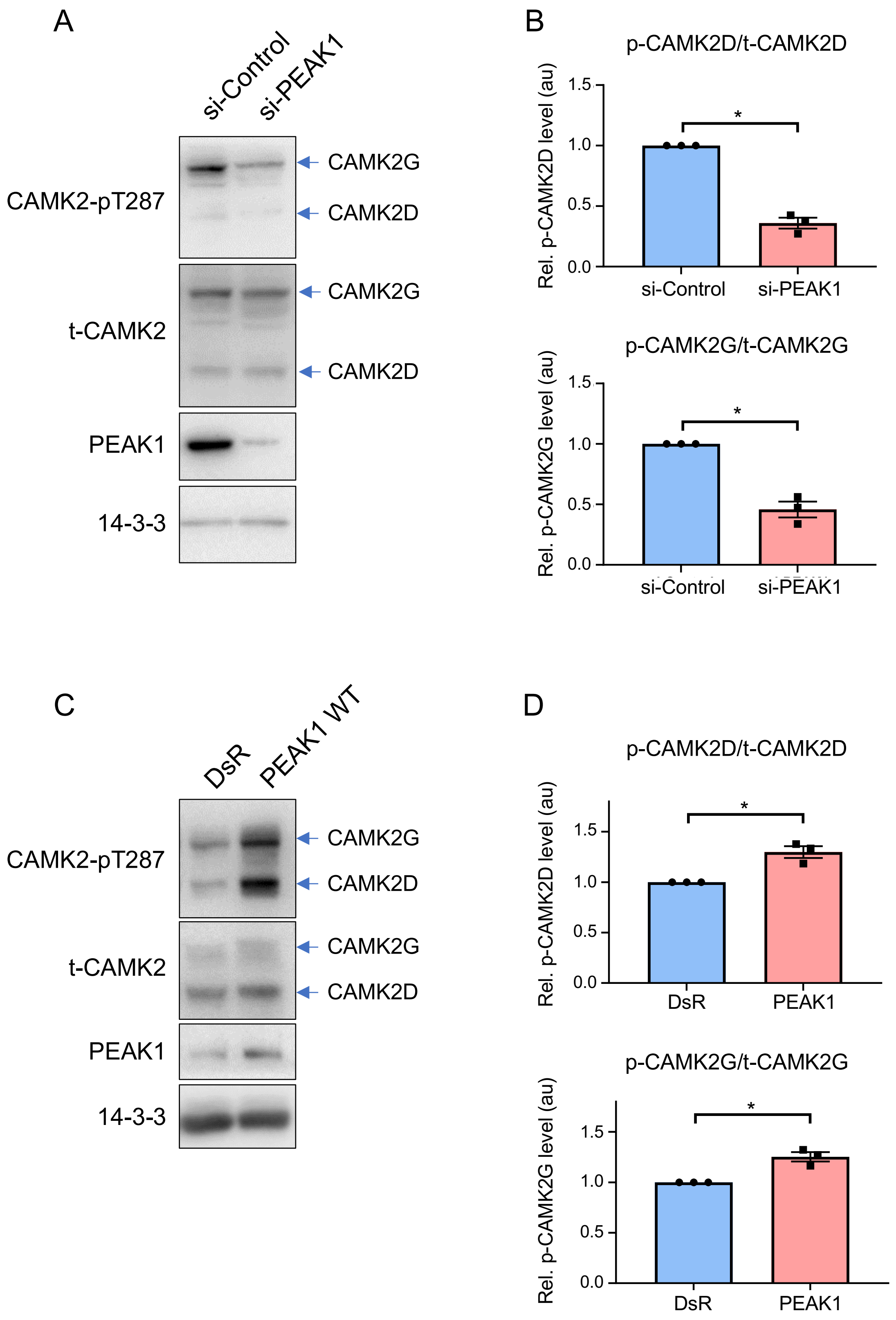


**Supp Figure 2. Manipulation of PEAK1 expression modulates CAMK2 activity in MDA-MB-468 cells.** PEAK1 was knocked down using siRNA (A-B) or overexpressed via transient transfection (C-D) in MDA-MB-468 cells and autonomous activation of CAMK2 determined by Western blotting as indicated. Histograms indicate CAMK2 T287 phosphorylation normalized for total CAMK2 expression, with 'au' indicating arbitrary units. * = $p < 0.05$ by ratio paired t-test.

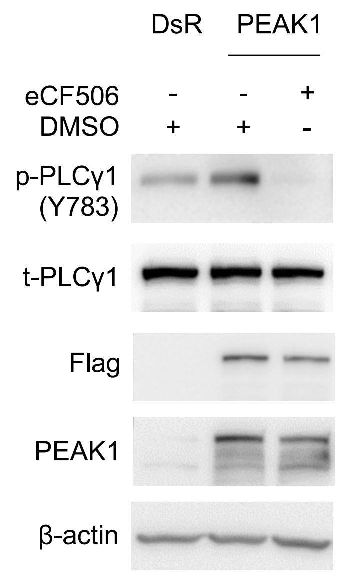


**Supp Figure 3. Effect of SFK inhibitor eCF506 on PLCγ1 Y783 phosphorylation.** MDA-MB-231_EcoR cells stably transduced with DsR empty vector or the corresponding PEAK1 construct were treated with DMSO or the SFK inhibitor eCF506 (250 nM) for 1 h. Cell lysates were then analysed by Western blotting as indicated. Data are representative of duplicate experiments.

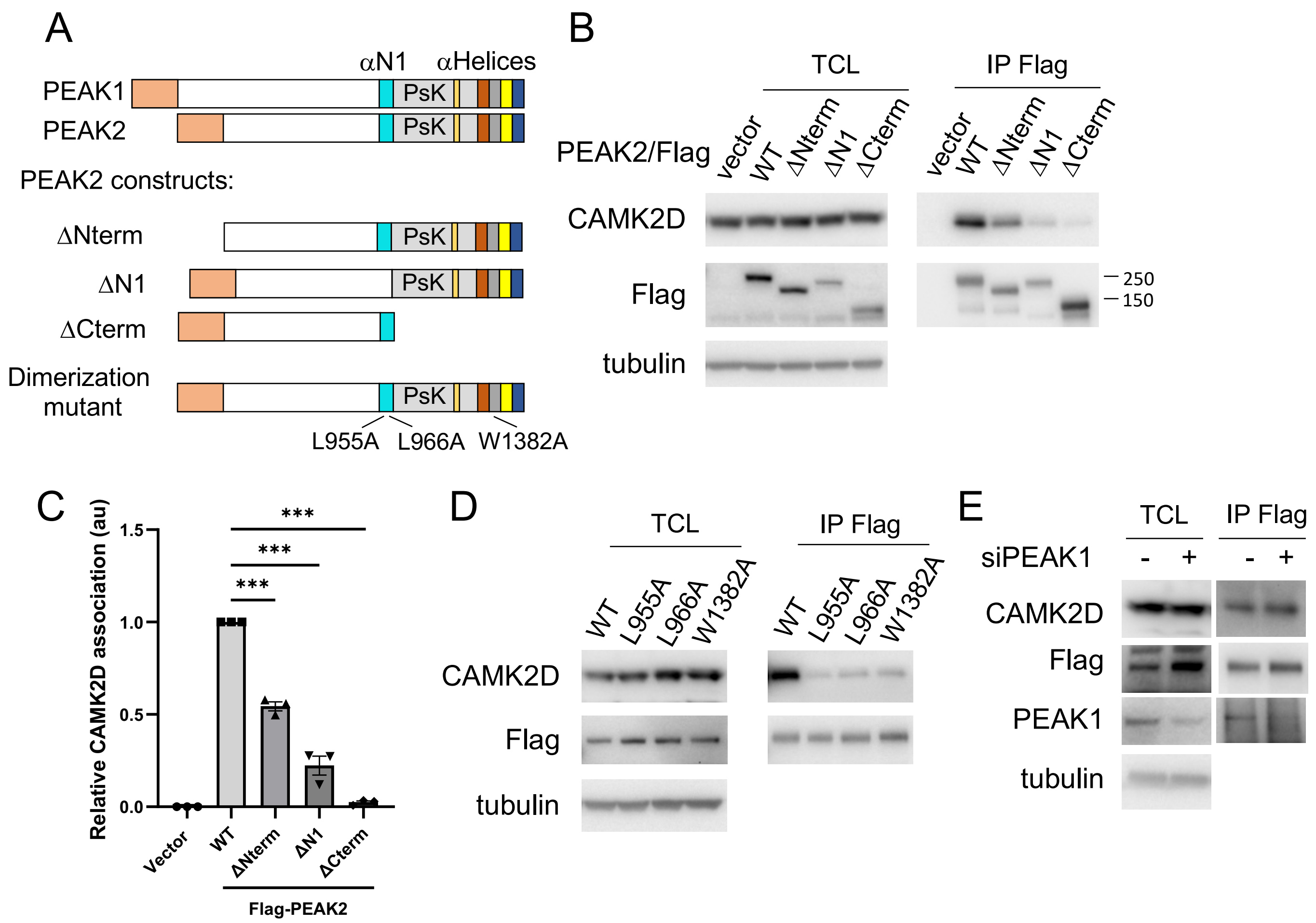


**Supp Figure 4. Determination of the structural requirements for CAMK2 interaction with PEAK2. A. Schematic structure of PEAK2 and various deletion and point mutants. B. Identification of PEAK2 regions required for association with CAMK2D.** Flag-tagged versions of the indicated PEAK2 proteins were transfected into HEK293T cells. Total cell lysates (TCL) and Flag immunoprecipitates (IPs) were subjected to Western blotting. **C. Quantification of CAMK2D association.** CAMK2D binding to PEAK2 was quantified and normalized for the PEAK2 level. Data are expressed relative to WT which was arbitrarily set at 1, with 'au' indicating arbitrary units. **D. Role of PEAK2 dimerization in regulating association with CAMK2D.** Flag-tagged versions of the indicated PEAK2 proteins were expressed in HEK293T cells. TCL and Flag IPs were subjected to Western blotting as indicated. **E. The association of PEAK2 with CAMK2D is PEAK1-independent.** HEK293T cells were transfected with a plasmid encoding Flag-tagged PEAK2 in the presence or absence of siRNA-mediated PEAK1 knockdown. TCL and Flag IPs were subjected to Western blotting as indicated. All data are representative of at least 2 independent experiments. Error bars in C represent the standard error of the mean from n=3 independent experiments. *** = $p < 0.001$ by one-way ANOVA with Dunnett's multiple comparisons test.

```
                    270        280        290        300        310
                  . |....|.... |....|.... |....|.... |....|.... |....|....
Human             Q PRFANFRANT LSPVRFFVDK KWNTIPLRNK SLQRICAVDY DDSYDEILNG
Mouse             Q PRFANFRANT LSPVRFFVSK KWNTIPLRNK SLQRICAVDY DDSYDEILNG
Rabbit            Q PRFANFRANT LSPVRFFVDK KWNTVPLRNK SLQRICAVDY DDSYDEILNG
Chicken           Q PRFANFRANT LSPVQFCVDK KWNTVPLRNK SLQRICAVDY DDSYDEILNG
Vampire bat       Q PRFANFRANT LSPVQFHVDK KWNTVPLRNK SLQRICAVDY DDSYDEILNG
Fox               Q PRFANFRANT LSPVRFFVDK KWNTVPLRNK SLQRICAVDY DDSYDEILNG
Squirrel          Q PRFANFRANT LSPVRFFVDK KWNTIPLRNK SLQRICAVDY DDSYDEILNG
Pangolin          Q PRFADFRADT LSPVRFSVDK KWNTVPLRNK SLQRICAVDY DDSYDEILNE
Hedgehog          Q PRFANFRANT LSPVRFFVGK KWNTVPLRNK SLQRICAVDY DDSYDEILNG
Gecko             Q PRFANLH-XT FSPFRFCIDK KWNTVPLRNK SLQRFCAVDY DDSYDEILNG
Komodo dragon     R PRFANFRANT LSPVRFCVDK KWNTVPLRNK SLQRFCAVDY DDSYDEILNG
Alligator         Q PRFANFRANT LSPVHFCADK KWNTVPLRNK SLQRICAVDY DDSYDEILNG
Tortoise          Q PRFANFRANT LSPVRFCVDK KWNTVPLRNK SLQRICAVDY DDSYDEILNG
Green sea turtle  Q PRFANFRANT LSPVRFYVDK KWNTVPLRNK SLQRICAVDY DDSYDEILNG
Tasmanian devil   Q PRFANFRANT LSPVRFFVGK KWNTVPLRNK SLQRICAVDY DDSYDEILHD
Echidna           Q PRFANFRANT LSPVQFSVGK KWNTVPLRNK SLQRICAVDY DDSYDEILNG
Platypus          Q PRFANFRANT LSPVQFSVGK KWNTVPLRNK SLQRICAVDY DDSYDEILNG
                  : ****:::  : :**. *   : ****:*:**: ****:***.* ********
                                                 R297  L301 R303
```

**Supp Figure 5. Sequence alignment of the PEAK1 CIM motif across diverse species.** The critical residues for CAMK2 interaction, R297, L301 and R303 are conserved across species and highlighted.

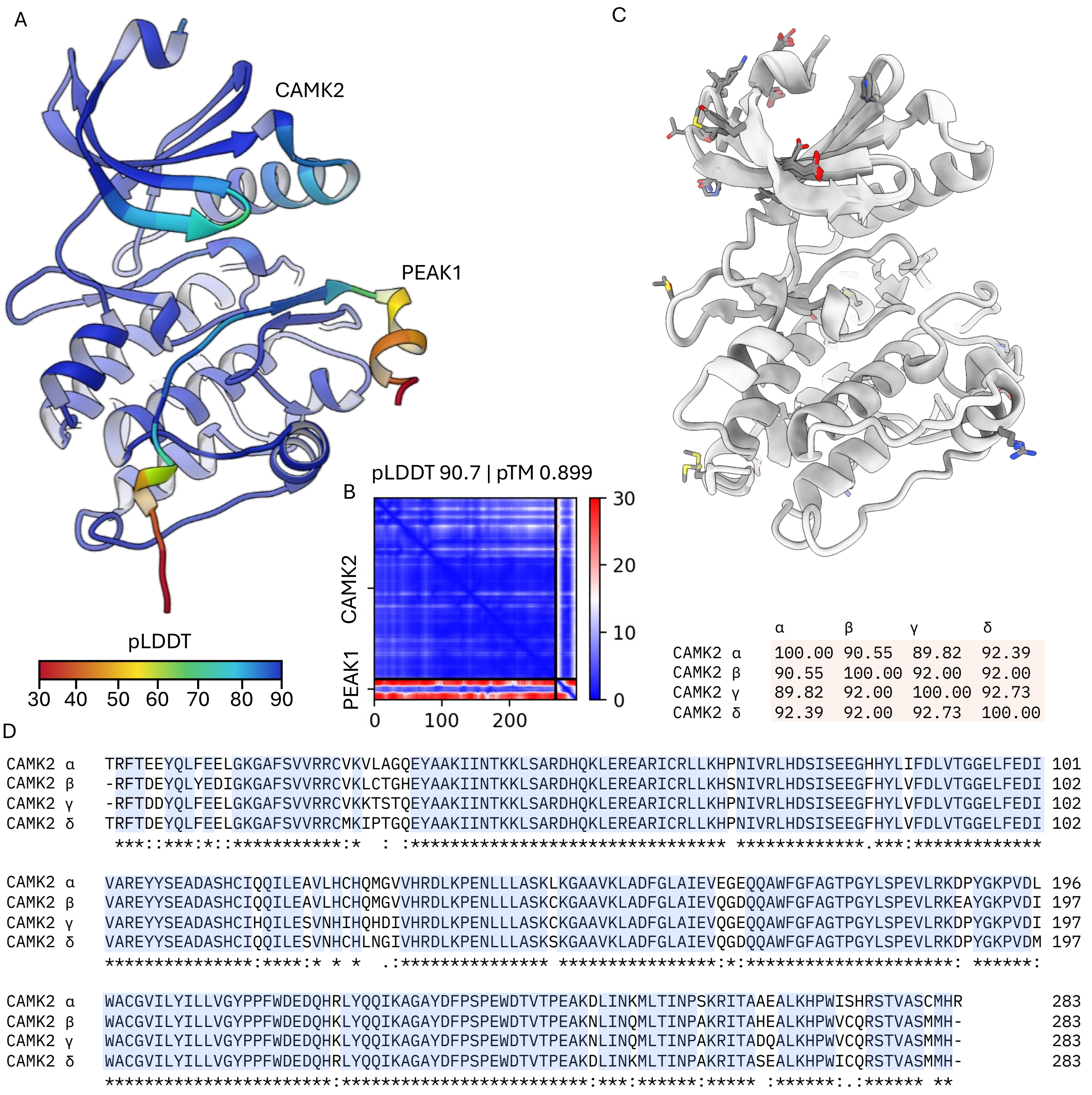


**Supp Figure 6. Comparative modelling analysis by AlphaFold (to accompany Figure 4C-F). A. Computationally predicted PEAK1(291-320)/CAMK2 model by AlphaFold (v3) coloured according to pLDDT score.** Overall pLDDT and pTM scores of 90.7 and 0.899 indicate high estimated confidence measured by per-residue local distance difference (LDDT) and residue-residue alignment error estimate respectively. **B. Predicted alignment error plot illustrating low overall estimated error between PEAK1(291-320) and CaMK2 (top right quadrant). C. Superimposition of CAMK2 isoforms.** Residues with less than 100% identity are shown as sticks. All other residues (identity 100%) are hidden for clarity. The residues that define the PEAK1(291-320)/CAMK2 interaction are conserved across all isoforms (α, β, γ, and δ). The identity matrix is shown below, with all isoforms having ~ 90% identity. **D**. Multiple sequence alignment of the domain region shown in (A) highlighting very high conservation between CAMK2 isoforms.

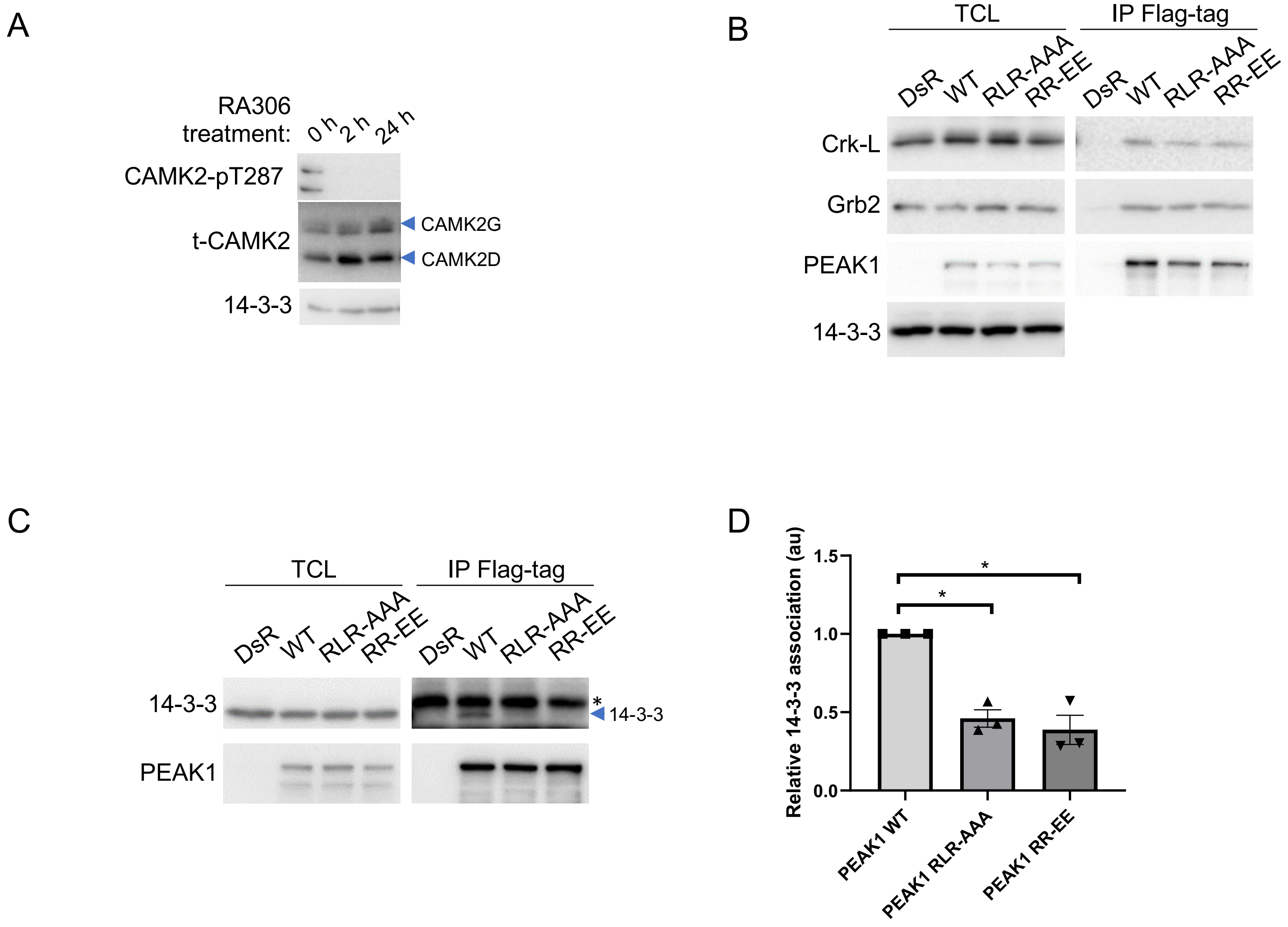


**Supp Figure 7. Role of CAMK2 in regulating PEAK1 phosphorylation and the PEAK1 interactome. A. Validation of CAMK2 inhibition using RA306.** MDA-MB-231 cells were treated with 1 µM RA306 for the specified times and then cell lysates were Western blotted as indicated. Data are representative of triplicate experiments. **B-D. Impact of CIM mutations on the PEAK1 interactome.** IPs of WT and mutant PEAK1 proteins from HEK293T cells were analysed by immunoblotting with the indicated antibodies. Results are representative of 3 independent experiments. The asterisk in C indicates a non-specific band. Histogram (D) indicates the association of 14-3-3 relative to WT PEAK1 (mean ± s.e.m., n = 3), with 'au' indicating arbitrary units. * = $p < 0.05$ by ratio t-test.

**A**

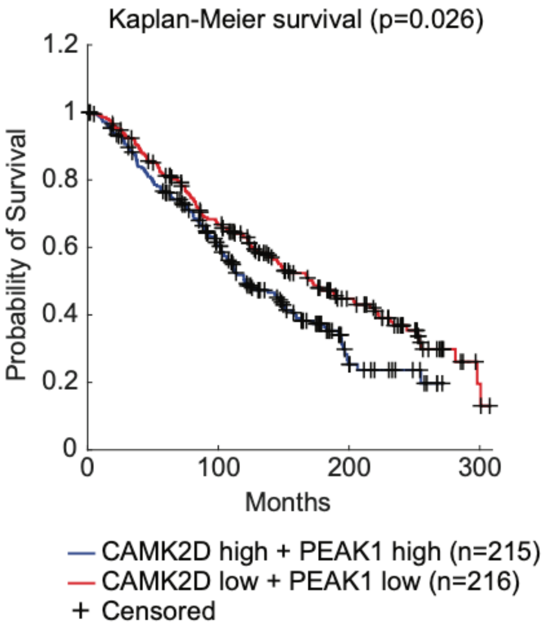


**B**

**Comparison**

| CAMK2D/PEAK1 | CAMK2D/PEAK1 | P-value |
|---|---|---|
| high/high | low/low | **0.026** |
| low/high | low/low | **0.8** |
| high/high | high/low | **0.051** |
| high/low | low/low | 0.71 |
| high/high | low/high | **0.026** |

**C**

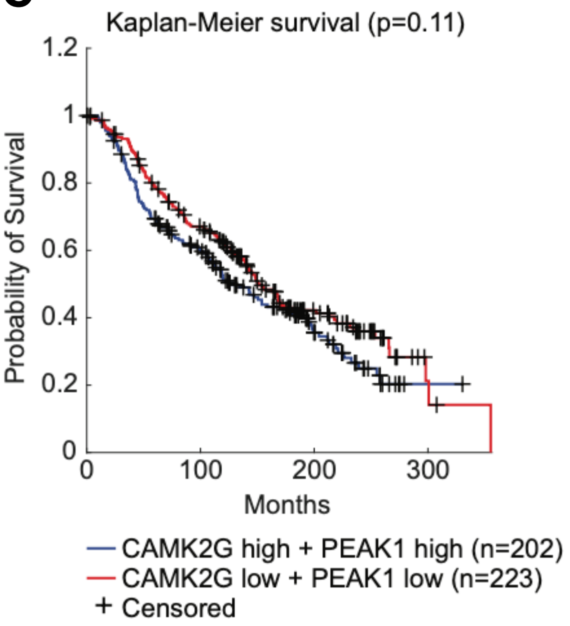


**D**

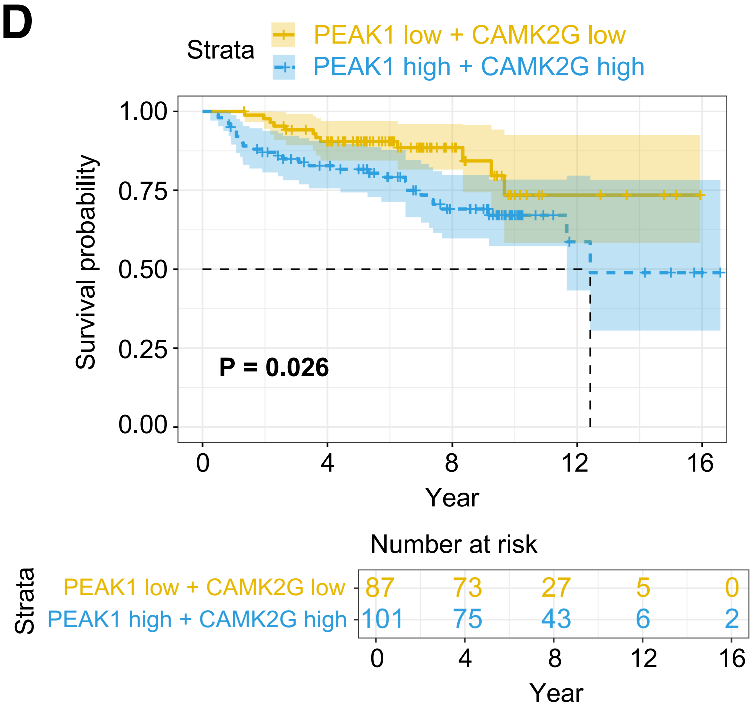


**Supp Figure 8. Association of PEAK1 and CAMK2 expression with breast cancer patient survival.** Gene expression, mutation profile and associated overall survival data from 2509 breast cancer patients were downloaded from the cBioPortal for Cancer Genomics portal (https://www.cbioportal.org/). Breast cancer patients were categorised into groups exhibiting different PEAK1/CAMK2 combined expression patterns. **A. Survival analyses comparing overall survival between PEAK1/CAMK2D both high and PEAK1/CAMK2D both low.** This was undertaken using a Log-rank test. **B. Summary table comparing overall survival differences between indicated PEAK1/CAMK2D combined expression patterns. C. Survival analyses comparing overall survival between PEAK1/CAMK2G both high and PEAK1/CAMK2G both low patient groups.** The Log-rank test statistics and survival curves were generated using Kaplan-Meier estimate and implemented using the Logrank package in MATLAB 2023a (with $p < 0.05$ considered significant). **D. Distant metastasis-free survival (DMFS) for the PEAK1/CAMK2G both high and PEAK1/CAMK2G both low patient groups.** DMFS was compared between the two subgroups using a log-rank test (with $p < 0.05$ considered significant). For cohort details, please refer to Methods.

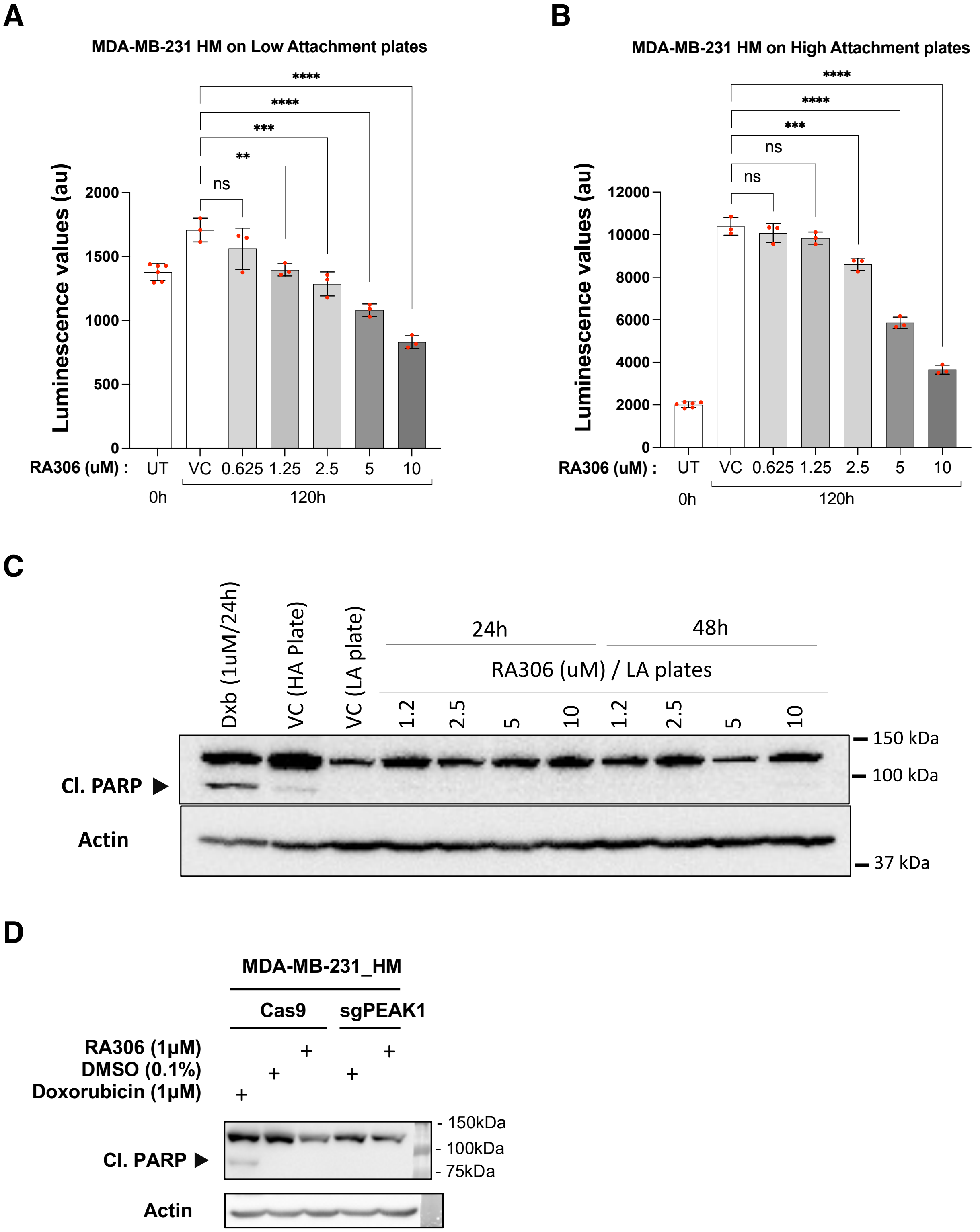


**Supp Figure 9. Effect of the CAMK2 inhibitor RA306 on cell proliferation and apoptosis. A-B, Effect on cell proliferation under low (A) and high (B) attachment conditions.** Cell viability assays were undertaken at the indicated time points, with data points indicating mean +/- standard deviation for triplicate wells. Data are representative of duplicate independent biological replicates. UT, untreated; VC, vehicle control (DMSO); au, arbitrary units. ns = not significant at p<0.05, ** = p <0.01, *** = p <0.001, **** = p

<0.0001, by one-way ANOVA with Dunnett's Multiple Comparison test. **C. Effect of RA306 on apoptosis under low attachment conditions.** MDA-MB-231 HM cell lysates were prepared at the indicated time after plating into low attachment plates and Western blotted as indicated. Doxorubicin (Dxb) treatment of cells on high attachment plates was used as a positive control for induction of cleaved (Cl) PARP. VC, vehicle control (DMSO) for 48h on high attachment (HA) and low attachment (LA) plates. Data are representative of duplicate biological replicates. **D. Effect of RA306 and PEAK1 gene knockout on apoptosis.** Control (Cas9) or PEAK1 gene knock-out (sgPEAK1) cells on HA plates were treated with vehicle control (DMSO) or RA306 for 24 h. Cell lysates were then Western blotted as indicated. Doxorubicin treatment was used as a positive control for induction of cleaved (Cl) PARP. Data are representative of duplicate biological replicates.

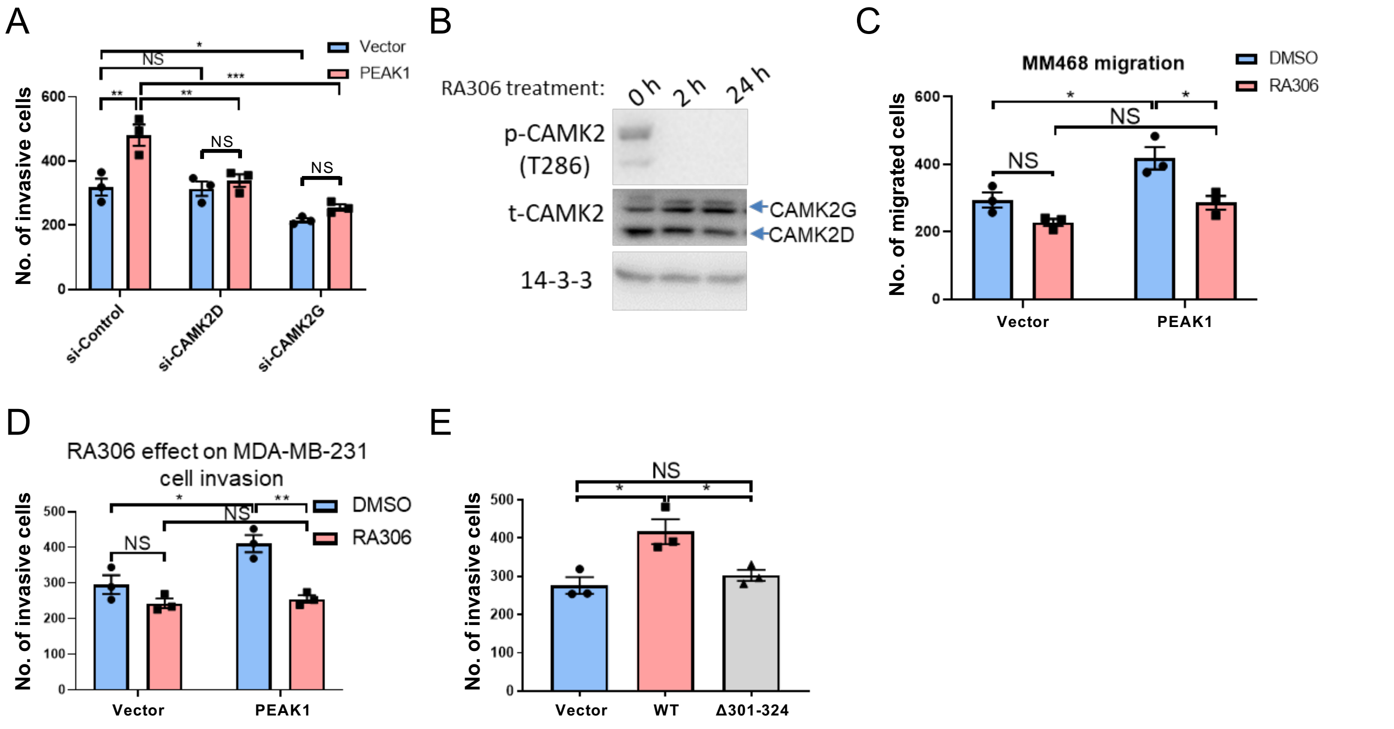


**Supp Figure 10. Role of CAMK2 in PEAK1-regulated biological endpoints in TNBC models. A. Role of CAMK2D/G in PEAK1-promoted MDA-MB-231 cell invasion.** MDA-MB-231 cells were transfected with a PEAK1 expression plasmid in the presence or absence of siRNA-mediated CAMK2D or CAMK2G knockdown. Cell lysates were validated by Western blotting in Figure 7A. Cells were subjected to a transwell invasion assay. **B-D. Pharmacological inhibition of CAMK2 using RA306 blocks PEAK1-promoted TNBC cell migration and invasion.** MDA-MB-468 cells were treated with RA306 for different times and cell lysates were then Western blotted as indicated (B). MDA-MB-468 cells were transfected with a PEAK1 plasmid and subject to transwell migration assays in the presence or absence of RA306 (C). MDA-MB-231 cells were transfected with a PEAK1 plasmid and subjected to transwell invasion assays in the presence or absence of RA306 (D). **E. Role of CAMK2 activation.** Plasmids expressing WT PEAK1 and the Δ301-324 mutant that cannot activate CAMK2 were transiently transfected into MDA-MB-231 cells, and cells were subjected to a transwell invasion assay. Error bars represent the standard error of the mean from n=3 independent assays. NS indicates $p > 0.05$, * = $p < 0.05$, ** = $p < 0.01$, *** = $p < 0.001$ by two-way ANOVA with Tukey's multiple comparisons test (A, C, D) or one-way ANOVA with Tukey's multiple comparisons test (E).

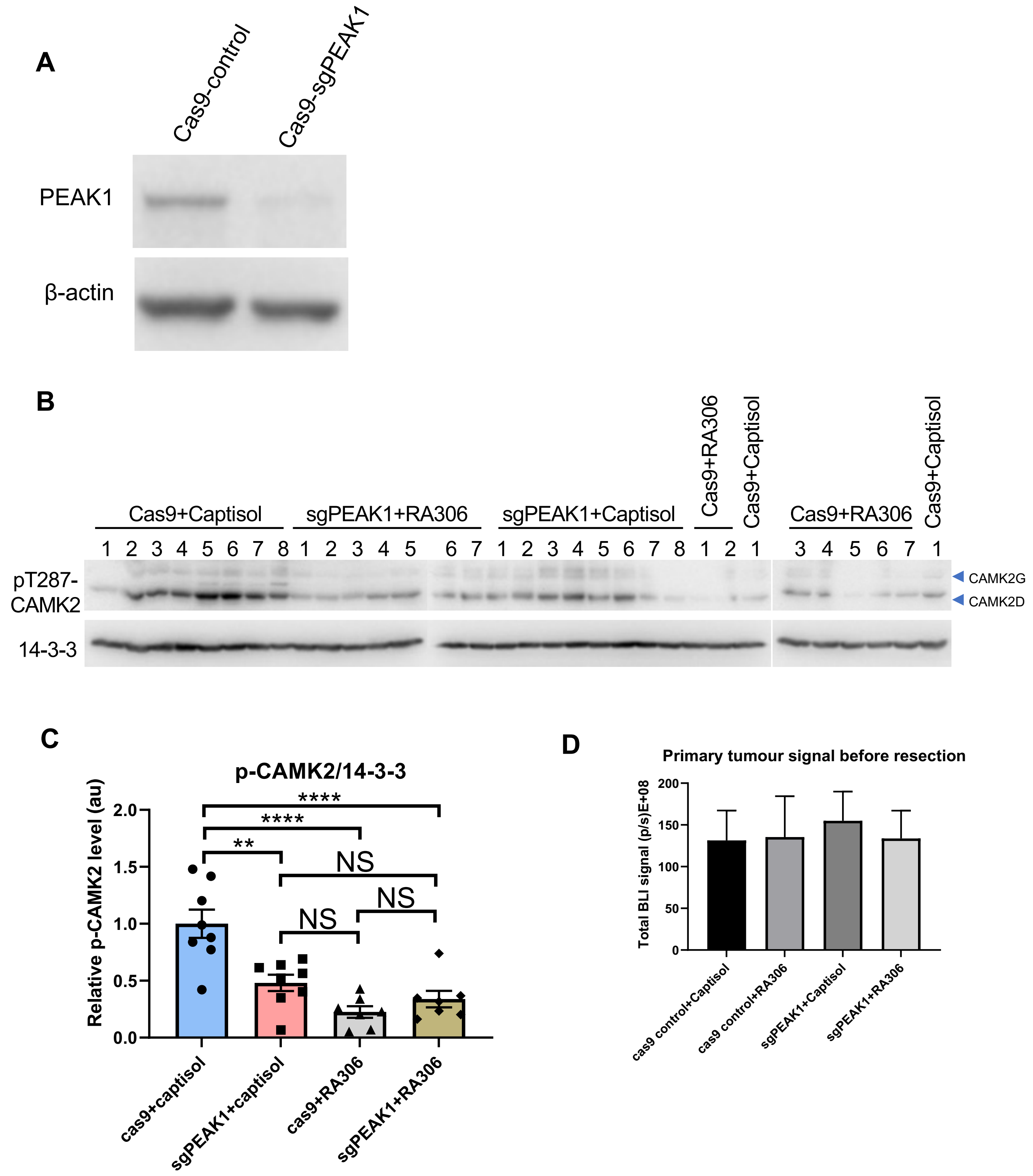


**Supp Figure 11. Genetic or pharmacological targeting of the PEAK1/CAMK2 axis reduces TNBC development *in vivo*. A. Validation of CRISPR-mediated PEAK1 knockout in MDA-MB-231_HM cells.** Cells were Western blotted as indicated. **B-C. CAMK2 activation in different xenograft treatment groups.** Tumour tissue from each group was harvested at 2 h after last RA306 treatment, homogenized and subjected to Western blotting (B) and quantification for combined CAMK2D and CAMK2G activation (C) as indicated. In C, data are expressed relative to Cas9-control/Captisol which was arbitrarily set at 1. Error bars represent the standard error of the mean. NS indicates $p > 0.05$, ** = $p < 0.01$, **** = $p < 0.0001$ by two-way ANOVA with Tukey's multiple comparisons test. **D. Primary tumour sizes prior to resection for metastasis experiment.** This accompanies Figure 7F. Data points are mean +/- SEM.
